# Assessing uncertainty in connective field estimations from resting state fMRI activity

**DOI:** 10.1101/2020.11.02.365452

**Authors:** Azzurra Invernizzi, Nicolas Gravel, Koen V. Haak, Remco J. Renken, Frans W. Cornelissen

## Abstract

Connective Field (CF) modeling estimates the local spatial integration between signals in distinct cortical visual field areas. As we have shown previously using 7T data, CF can reveal the visuotopic organization of visual cortical areas even when applied to BOLD activity recorded in the absence of external stimulation. This indicates that CF modeling can be used to evaluate cortical processing in participants in which the visual input may be compromised. Furthermore, by using Bayesian CF modelling it is possible to estimate the co-variability of the parameter estimates and therefore, apply CF modeling to single cases. However, no previous studies evaluated the (Bayesian) CF model using 3T resting-state fMRI data, although this is important since 3T scanners are much more abundant and more often used in clinical research than 7T ones. In this study, we investigate whether it is possible to obtain meaningful CF estimates from 3T resting state (RS) fMRI data. To do so, we applied the standard and Bayesian CF modeling approaches on two RS scans interleaved by the acquisition of visual stimulation in 12 healthy participants.

Our results show that both approaches reveal good agreement between RS- and visual field (VF)-based maps. Moreover, the 3T observations were similar to those previously reported at 7T. In addition, to quantify the uncertainty associated with each estimate in both RS and VF data, we applied our Bayesian CF framework to provide the underlying marginal distribution of the CF parameters. Finally, we show how an additional CF parameter, *beta*, can be used as a data-driven threshold on the RS data to further improve CF estimates. We conclude that Bayesian CF modeling can characterize local functional connectivity between visual cortical areas from RS data at 3T. In particular, we expect the ability to assess parameter uncertainty in individual participants will be important for future clinical studies.

**Highlights:**

- Local functional connectivity between visual cortical areas can be estimated from RS-fMRI data at 3T using both standard CF and Bayesian CF modelling.
- Bayesian CF modelling quantifies the model uncertainty associated with each CF parameter on RS and VF data, important in particular for future studies on clinical populations.
- 3T observations were qualitatively similar to those previously reported at 7T.

## 1. Introduction

Spontaneous blood-oxygen level dependent (BOLD) fluctuations have been used to study the intrinsic functional connectivity of the brain. In 1995, *Biswal and colleagues* observed, for the first time, the presence of bilateral spatial integration, coherent activity and functional connectivity between distant homotopic brain areas, even in the absence of a task (Biswal *et al.*, 1995). Ever since, resting-state fMRI (RS-fMRI o RS) has played a key role in understanding the temporal and spatial interactions of interconnected brain regions. In parallel, various fMRI data-analysis tools have been developed with the aim to describe and estimate functional and neuroanatomical organization. One of these methods is connective field (CF) modeling (Haak *et al.*, 2013a). Even though this approach is stimulus agnostic by design, thus far it has been primarily developed and applied in vision research. CF modeling allows to characterize the response of a population of neurons in the cortex in terms of the activity in another region of the cortex. It translates the concept of the receptive field into the domain of connectivity by assessing the spatial dependency between signals in distinct cortical visual field regions (Haak *et al.*, 2013a). This type of receptive field is also known as the cortico-cortical population receptive field (cc-pRF). A previous study by *Gravel et al.* showed that CFs, estimated from RS-fMRI data recorded at a high magnetic field (7T), reflect the visuotopic organization of early visual cortical maps (Gravel *et al.*, 2014). This indicates that even in the absence of any visual stimulation, CF modeling is able to describe the activity of voxels in a target region (e.g. V2 or V3) as a function of the aggregate activity in a source cortical visual area (e.g. V1).

While these previous results were obtained at 7T and in healthy participants, 3T scanners are much more common, and generally preferred for whole-brain analyses in patient studies (Kolk *et al.*, 2013; Polimeni and Uludağ, 2018). Therefore, if RS data recorded at 3T can provide sufficient sensitivity to estimate the spatial integration and connectivity of BOLD signals in distinct regions of the early visual cortex (Gravel *et al.*, 2020), this would open up the CF modeling approach to clinical studies performed at 3T and in individual cases. Amongst others, this would give the advantage that plasticity of visual cortical areas could be studied without a dependence on actual visual stimulation. This is important, as in ophthalmic and neurological patients visual input can already be disrupted, potentially resulting in spurious plasticity (Baseler, Morland and Wandell, 1999; Azzopardi and Cowey, 2001; Haak *et al.*, 2013a).

To demonstrate feasibility of the CF approach at 3T, we will compare newly obtained 3T results with 7T results previously obtained in a different cohort of healthy subjects. Moreover, in order to assess the suitability of 3T data in unique patient cases, we will look beyond the classical variance explained as an indicator of modeling performance and we will directly assess the uncertainty of model parameter estimates using a Bayesian approach. These parameters are available to us by applying our recently developed Bayesian framework for the CF model (Bayesian CF, Invernizzi *et al.*, 2020). In particular, this approach allows to estimate the variability for each CF parameter estimate such as CF size and beta. Moreover, when using our new Bayesian CF framework, we can obtain a data-driven threshold in order to select relevant voxels for both RS-fMRI and visual field mapping (VFM) data.

In order to compare our results to those obtained previously in Gravel *et al.*, we applied both the standard CF estimation and the novel Bayesian approach to RS and VFM data acquired at 3T and subsequently, we compared the CF maps and parameters obtained using different CF models. Additionally, we assessed test-retest reliability between the two runs of RS data.

To preview our results, we found a good agreement between RS- and visual field (VF) - based maps obtained with both the standard and Bayesian CF approach. Moreover, most observations were similar to those previously observed for 7T data. This implies that local functional connectivity between visual cortical areas during RS can be estimated at 3T. No significant differences were found between the two runs of RS data. Furthermore, we showed how the parameter uncertainty can be used to assess the variability of parameters in RS-fMRI BOLD fluctuations. Therefore, the Bayesian CF approach presented here provides an interpretable and independent measure of uncertainty in both VFM- and RS-based data. Finally, we show that the novel retained CF parameter, *beta*, can serve as a sensitive threshold for the selection of voxels and improve the reliability of estimates.

## 2. Methods

### 2.1 Participants

Twelve healthy female participants (mean age 22 years, s.d. = 1.8 years) with normal or corrected-to-normal vision and without a history of neurological disease were included. These data were already used in previous projects (Halbertsma, Haak and Cornelissen, 2019; Invernizzi *et al.*, 2020). The ethics board of the University Medical Center Groningen (UMCG) approved the study protocol. All participants provided written informed consent. The study followed the tenets of the Declaration of Helsinki.

### 2.2 Stimuli presentation and description

The visual stimuli were displayed on a MR compatible screen located at the head-end of the MRI scanner with a viewing distance of 118 cm. The participant viewed the complete screen through a mirror placed at 11 cm from the eyes supported by the 32-channel SENSE head coil. Screen size was 36 × 23 degrees of visual angle and the distance from the participant’s eyes to the screen was approximately 75 cm. Stimuli were generated and displayed using the Psychtoolbox (https://github.com/Psychtoolbox-3/Psychtoolbox-3/) and VISTADISP toolbox (VISTA Lab, Stanford University), both MatLab based (Brainard, 1997; Pelli, 1997). The stimulus consisted of drifting bar apertures (of 10.2 deg radius) with a high contrast checkerboard texture on a grey (mean luminance) background. A sequence of eight different bar apertures with four different bar orientations (horizontal, vertical and diagonal orientations), two opposite motion directions and four periods of mean-luminance presentations completed the stimulus presentation that lasted 192 second. To maintain stable fixation, participants were instructed to focus on a small colored dot present in the center of the screen and press a button as soon as the dot changed color. The complete visual field mapping paradigm was presented to the participant six times, during six separate scans.

### 2.3 Resting state

During the RS-fMRI scans, the stimuli were replaced by a black monitor and the lights in the scanning room were turned off. All participants were instructed to keep their eyes closed, remain as still as possible, not to fall asleep and try not to think of anything in particular.

### 2.4 Data acquisition

MRI and fMRI data were obtained using a 3T Philips Intera MRI scanner (Philips, the Netherlands), with a 32-channel head coil. For each subject, a high-resolution T1-weighted three-dimensional structural scan was acquired (TR = 9.00ms, TE = 3.5ms, flip-angle = 8, acquisition matrix = 251*251*170mm, field of view = 256×170×232, voxel size = 1×1×1mm). Then, functional T2*-weighted, 2D echo planar images were obtained (TR = 1500ms, TE = 30ms,field of view = 190×190×50 mm, voxel resolution of 2.5×2.5×2.5). For the functional data, the same protocol was used for collecting both RS and VFM sequences. Only the TR parameter was modified to 2000ms for the RS-fMRI data. The functional scans were acquired in the following order: (1) a RS-fMRI scan (RS1) lasted 370s with a total of 340 volumes; (2) six VFM functional scans were collected, where each scan lasted for 192s with a total of 136 volumes; (3) finally, a second RS-fMRI scan (RS2) with the same characteristic of RS1 (duration of 370s with 340 volumes) was collected. Prior to the first VFM scan, a short T1-weighted anatomical scan with the same field of view chosen for the functional scans were acquired and used for obtaining a better co-registration between functional and anatomical volume.

### 2.5 Data analysis

Preprocessing and standard CF analysis of fMRI data were done using ITKGray (http://www.itk.org), Freesurfer (Fischl, 2012) and mrVista (VISTASOFT) toolbox from Stanford University (http://www.white.stanford.edu). The Bayesian pRF and CF approaches were developed and implemented in MatLab 2016b (The Mathworks Inc., Natick, Massachusetts). The code for the MCMC pRF and CF frameworks will be made available via the website www.visualneuroscience.nl.

For each subject, the structural scan was aligned in a common space defined using the anterior commissure-posterior commissure line (AC-PC line) as reference. Next, grey and white matter were automatically segmented using Freesurfer and manually adjusted using ITKGray (http://itk.org), in order to minimize segmentation errors. Then, all functional data were pre-processed using mrVista toolbox. For both RS and retinotopy data the following steps are applied. First, head motion within and between scans were corrected by using robust multiresolution alignment of MRI brain volumes (Nestares and Heeger, 2000) an alignment of functional data into anatomical space and an interpolation of functional data with segmented anatomical grey and white matter. For RS-fMRI data, a few additional steps were applied. First, spatial smoothing was applied (6 millimeters FWHM) then the denoising step based on ICA-AROMA was applied to remove the noise components from the unsmoothed RS-fMRI data (Pruim *et al.*, 2015). Then, an additional band-pass filter with high-pass discrete cosine transform with cut-off frequency of 0.01 Hz and a low-pass 4th order Butterworth filter with cut-off frequency of 0.1 Hz were applied to RS data.

#### 2.5.2 Bayes population receptive field mapping applied to VFM

Retinotopy scans were analyzed using a Bayes population receptive field (pRF) framework. For a detailed account see *Prabhakaran et colleagues,* which uses a Markov Chain Monte Carlo (MCMC) approach to sample the parameter space for the pRF mapping. Following the nomenclature of (Dumoulin and Wandell, 2008; Zeidman *et al.*, 2018; Carvalho *et al.*, 2020; Prabhakaran *et al.*, 2020), we defined 2D symmetrical Gaussian kernel centered at (*x*_0_, *y*_0_) with width defined as the standard deviationσ, to define the pRF model. The best model fit was projected onto a smoothed 3D mesh of the cortex. Based on the obtained parameter-values, visual areas are outlined (V1, V2, V3, hV4, LO1 and LO2) to act as source (V1) or target region (all other) for subsequent RS analysis.

#### 2.5.3 Standard connective field mapping of RS data

In the standard CF model, the optimal CF parameters (CF position and CF size, which define the 2D symmetric Gaussian kernel) were estimated based on a procedure that fitted the time-series for each location in the target region (e.g. V2 or V3) using a linear combination of the time-series in the source region (e.g. V1; Haak *et al.*, 2013b). The best fitting models are retained and projected on a smoothed 3D mesh. The CF parameters associated with the best fitting model are converted from cortical units (cortical position) into visual field units (eccentricity and polar angle). This is done by inferring the pRF properties — obtained via the Bayesian pRF method (Prabhakaran *et al.*, 2020) of the center voxel in the source region for each target location (Haak *et al.*, 2013b).

#### 2.5.4 Bayesian connective field mapping

Similar to the Bayes pRF, the Bayesian CF framework uses a Markov Chain Monte Carlo (MCMC) approach to sample the source region efficiently. Again we used a 2D symmetric Gaussian kernel to predict the time series of the target regions. As in the standard CF modeling, the eccentricity and polar angle values associated with the CF centers are inferred from a pRF mapping. For sake of completeness, the complete fitting procedure of Bayesian CF model (option B) is described in the supplementary material (Invernizzi *et al.*, 2020).

For each subject, standard and Bayesian-CF models were estimated for both VFM and RS data. Target and source regions definitions were based on the Bayes pRF analysis. For both Bayes pRF and CF models, a total of 15000 iterations were computed, where the first 10% of iterations were discarded for the burn-in process (Chib, 2011; Liu, Nordman and Meeker, 2016) and the posterior probability distributions were estimated on the remaining samples.

#### 2.5.5 Spatial analysis

We used Pearson and circular correlations to compare and assess the topographic organization of eccentricity and polar angle, respectively, in both standard CF and Bayesian CF maps obtained on RS and VFM data. The same type of correlations and permutation tests were used to evaluate similarities in eccentricity and polar angle maps between the two RS-fMRI scans obtained with standard CF and Bayesian CF models. P-values below 0.05 were considered statistically significant. Moreover, to compare the relation between CF size and eccentricity RS-based, we binned the eccentricity at 1 degree intervals and applied a linear fit over the mean per bin. A confidence interval (CI) of the fit was defined by applying a bootstrap technique 1000 times.

For the all spatial analyses, only voxels for which the best-fitting CF model explained more than 15% of the time-series variance in the standard CF and eccentricity which is < 1deg and > 7deg were included.

#### 2.5.6 Bayesian analysis

Based on a quantile analysis of the posterior distribution (Invernizzi *et al.*, 2020), we computed a voxel-wise uncertainty measure for each CF parameter by subtracting the upper (*Q* _3_) and lower (*Q*_1_) quantile of the posterior distribution. The estimated uncertainty was computed for both RS and VFM data and then projected onto a smoothed 3D mesh of the cortex. We repeated the same procedure for each CF parameter.

#### 2.5.7 Beta threshold

Following the procedure reported by Invernizzi et al., we test if *beta* – the scaling amplitude of the predictor to the amplitude of the measured signal – can serve as data-driven threshold for RS-data. As a proxy distribution for the null hypothesis (i.e. no correlation between source and target region), one surrogate BOLD time series was calculated for each voxel (Schreiber and Schmitz, 1996; Räth and Monetti, 2009; Lancaster et al., 2018). A surrogate time series was generated using the iterative amplitude adjusted Fourier transform method (iAAFT) (Schreiber and Schmitz, 1996; Räth and Monetti, 2009). Then, the Bayesian CF model was fitted using this surrogate to real time series of the target region which were unchanged. Based on the best fit obtained in the MCMC iterations of the surrogate *beta*-estimate, we calculated a familywise error (FWE) corrected *beta-*threshold for all the voxels in the target region. To obtain the FWE *beta-*threshold, we used the 95th percentile of the null distribution as threshold. Finally, we compared the voxel selection at the single participant level using VE and the FWE *beta-*threshold approach.

## 3. Results

The CF maps obtained from RS-based data for eccentricity, polar angle and CF size were qualitatively comparable for the standard and Bayesian CF models. In contrast to the VFM data, the relation with CF size and eccentricity in RS data does not increase with visual hierarchy. Again, the same behavior was observed using both methods. A significant difference in eccentricity and variance explained (VE) parameters was found in VFM-*versus* RS1 - based data, while a significant difference in CF size was noticed in VFM - *versus* RS2-based data. We observed a higher value of variance explained (VE) of RS2 compared to RS1. No statistically significant difference was found between the two RS scans for any other CF parameter. We estimated the uncertainty for the Bayesian CF parameters (CF size and beta). An higher uncertainty from the CF parameters was observed from both RS scans compared to VFM data and between RS2 and RS1 scans. Finally, we showed how to use a new threshold based on the effect size of the model both in the presence and absence of visual stimuli.

### 3.1 CF models based on RS-fMRI data

We used V1 as source region while V2, V3, hV4, LO1 and LO2 as target to derive CF maps projected on a smoothed 3D mesh on a single subject level (Figure 1). Such maps were created using both standard CF and Bayesian CF models (Figure 1, panel B, C, D, and E). Topographical maps for eccentricity, polar angle and CF size were comparable for both CF models using RS data (Figure 1). We used the VFM-based maps as reference (Figure 1, Panel A) as these maps show a clear visuotopic organization for all the CF parameters estimated. Then the same parameters are plotted for all RS scans (Figure 1 - Panel B and D: RS-based derived maps using standard CF model; Panel C and E: RS-based derived maps using Bayesian CF model). To quantify the visuotopic organization of the resulting RS-based CF maps, we correlated the CF-derived eccentricity and polar angle to VFM-derived CF maps (Table 1). The best agreement was found for V1 > V2 areas using both models.

**Table 1.** Group level correlation between CF maps derived from visual field and resting state data obtained using Bayes pRF and CF modeling. To estimate the level of agreement between pRF and CF maps obtained by using VFM and RS data, we computed the correlations coefficients between the eccentricity (*rho*) and polar angle (*theta*) parameters obtained using standard CF and those derived using Bayesian CF models to the pRF *rho* and *theta* (gold standard). VFM values have been included as reference.

| ROIs | VFM |  | RS1 |  | RS2 |  | VFM - MCMC |  | RS1 - MCMC |  | RS2 - MCMC |  |
| --- | --- | --- | --- | --- | --- | --- | --- | --- | --- | --- | --- | --- |
|  | *rho* | *theta* | *rho* | *theta* | *rho* | *theta* | *rho* | *theta* | *rho* | *theta* | *rho* | *theta* |
| V1 --> V2 | 0.8619 | 0.7328 | 0.4716 | 0.5247 | 0.5397 | 0.5400 | 0.8576 | 0.7359 | 0.4073 | 0.5411 | 0.5397 | 0.5514 |
| V1 --> V3 | 0.8094 | 0.7051 | 0.2294 | 0.3141 | 0.2857 | 0.0583 | 0.8075 | 0.7010 | 0.2345 | 0.3371 | 0.2613 | 0.0324 |
| V1 -> hV4 | 0.7759 | 0.6851 | 0.1217 | -0.0719 | 0.2482 | 0.2330 | 0.7698 | 0.6787 | 0.0941 | -0.0362 | 0.2325 | 0.1506 |
| V1 -> LO1 | 0.7740 | 0.6600 | 0.0768 | 0.4923 | 0.1465 | 0.3115 | 0.7627 | 0.6645 | 0.0626 | 0.5004 | 0.1567 | 0.3744 |
| V1 -> LO2 | 0.5726 | 0.6292 | -0.0392 | 0.2758 | 0.1288 | 0.3050 | 0.5800 | 0.6064 | -0.0672 | 0.2373 | 0.0942 | 0.2999 |

**Figure 1.**
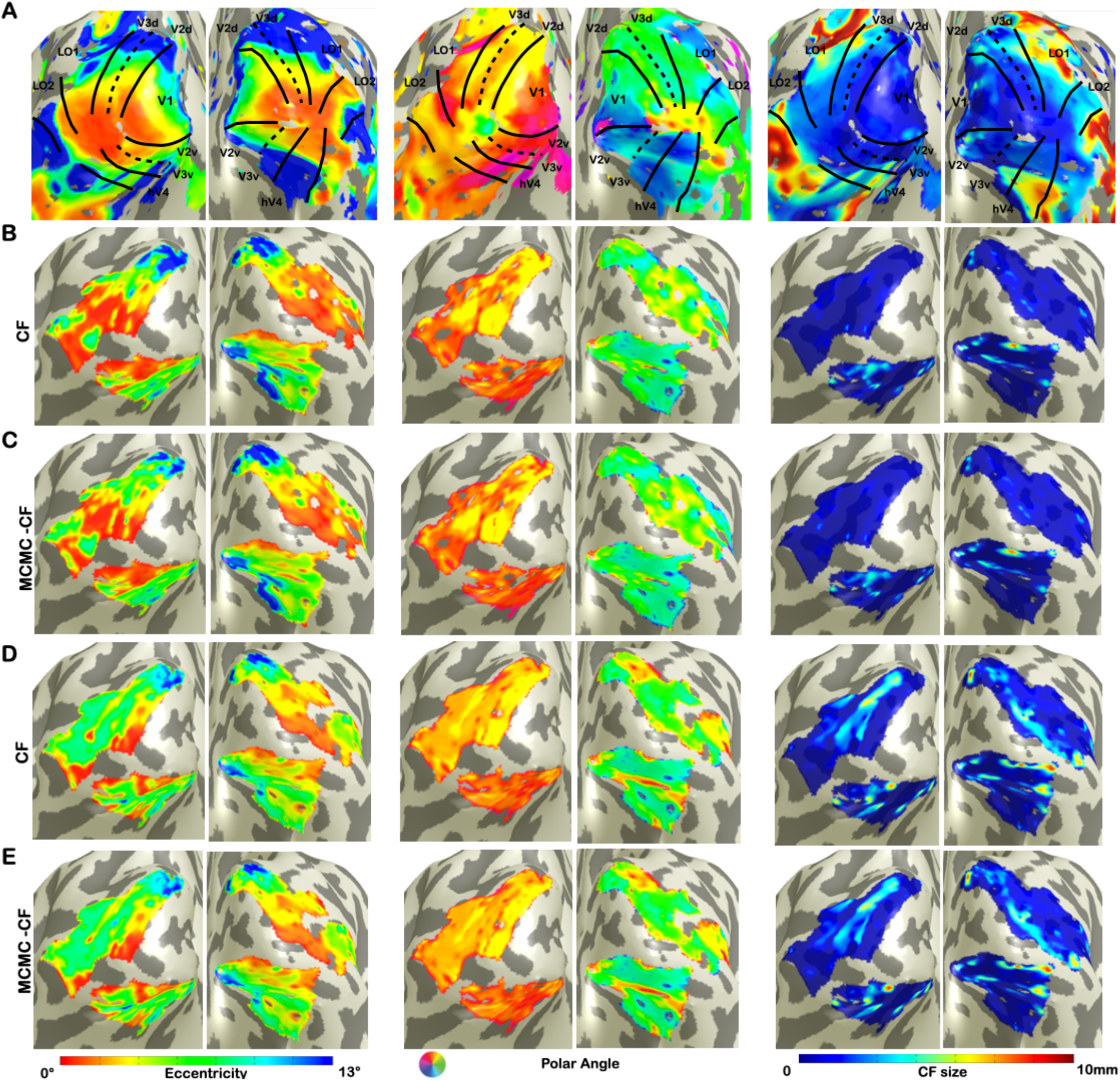
Visualization of CF maps of denoised data for a single subject. From left to right: eccentricity, polar angle and CF size. Panel A corresponds to VFM derived estimates. Panels B and D show parameter estimates for the first RS run using standard CF and Bayesian CF models, respectively. Panels C and E show parameter estimates for the second RS run using standard CF and Bayesian CF models, respectively.

To check the possible influence of the denoise procedure applied to RS data, the same quantification analysis was computed on non-denoised RS data. Similar results were observed indicating that the ICA-AROMA denoise procedure on RS-fMRI data did not influence the final CF outcomes (Supplementary material: Figure S1 and Table S1).

### 3.2 Assessing uncertainty in RS-fMRI data

In order to estimate a voxel-wise uncertainty value associated to each CF parameter, we computed a quantile analysis of the posterior distribution and projected on a smoothed 3D mesh for a single subject (Figure 2). We used V1 as source regions while V2, V3, hV4, LO1 and LO2 as target regions; VFM-based CF maps were used as reference (Fig. 2*A*). An increased uncertainty in beta estimate in RS1- and RS2-based CF maps was observed compared to VFM-based CF maps but not for CF size. Interestingly, no clear uncertainty-related visuotopic organization was found either for VFM or RS data.

**Figure 2.**
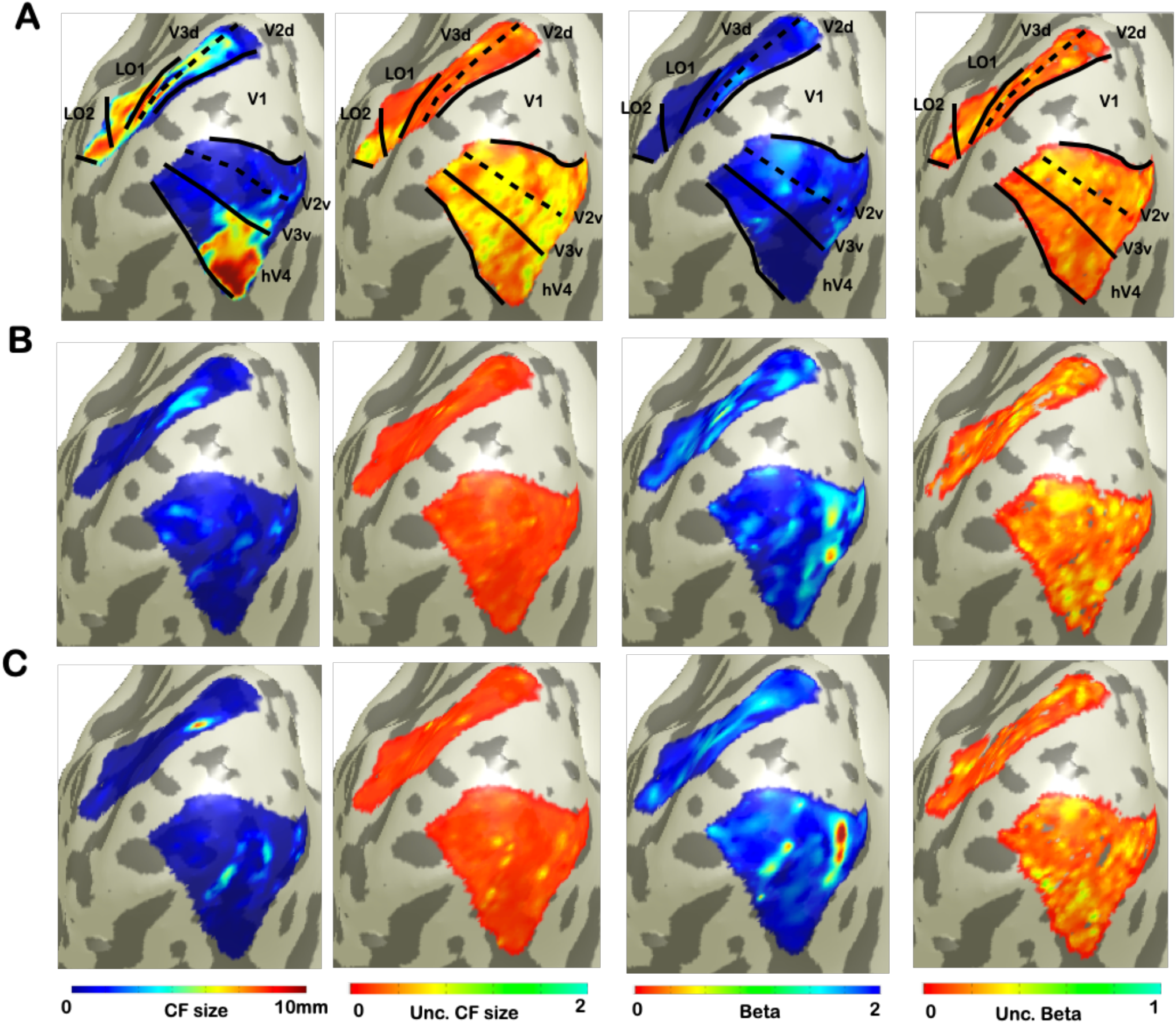
Visualization of uncertainty for CF parameter at single subject. From left to right: CF size, uncertainty of CF size, beta, and uncertainty of beta. Panel A corresponds to VFM derived estimates. While, bottom panels B and C show the parameters and uncertainty estimates for each RS scans.

As reported in *Invernizzi et al.* for VFM data, a weak correlation exists between beta, sigma parameter estimates and their respective uncertainties obtained on RS-data (Table 2).

**Table 2.** Dependency between MCMC CF parameters and uncertainties at voxel level for both RS scan. Cross-correlations were computed between the estimated MCMC CF parameters and the uncertainty derived from them. Only the CF parameters directly estimated using the model (CF size and beta) are included in this analysis.

|  | V1 > V2 |  | V1 > V3 |  | V1 > hV4 |  | V1 > LO1 |  | V1 > LO2 |  |
| --- | --- | --- | --- | --- | --- | --- | --- | --- | --- | --- |
| RS1 | CF size | Beta | CF size | Beta | CF size | Beta | CF size | Beta | CF size | Beta |
| Unc. CF size | 0.06 | -0.01 | 0.11 | 0.04 | 0.1 | 0.07 | 0.15 | 0.1 | 0.1 | 0.09 |
| Unc. Beta | -0.03 | 0.01 | -0.01 | 0.01 | -0.04 | 0.01 | -0.03 | -0.12 | -0.01 | -0.01 |
| RS2 | CF size | Beta | CF size | Beta | CF size | Beta | CF size | Beta | CF size | Beta |
| Unc. CF size | 0.08 | -0.08 | 0.06 | -0.05 | 0.04 | -0.02 | 0.03 | 0.02 | -0.02 | -0.06 |
| Unc. Beta | 0.01 | -0.01 | 0.01 | -0.02 | 0.01 | -0.03 | -0.03 | -0.04 | -0.02 | 0.04 |

### 3.3 Bayesian CF thresholding application

To evaluate the goodness of the corrected beta-thresholding method in the voxel selection on RS data, we compared the model VE, each CF parameter and the uncertainty associated, respectively (Figure 4, CF size and Figure 2S, beta parameter).Both thresholds: VE is higher than 15% and the FWE corrected effect size (>95% Figure 4, Panel A; for more details, see Invernizzi *et al.*, 2020) are indicated. Based on a direct comparison of FWE *beta*-corrected threshold (CI) to the standard VE of the model (Figure 4, Panel B and C), the 95% FWE CI-based threshold proved to be more conservative. Note that it is not straightforward to identify a point at which the two threshold definitions will be equivalent.

**Figure 4.**
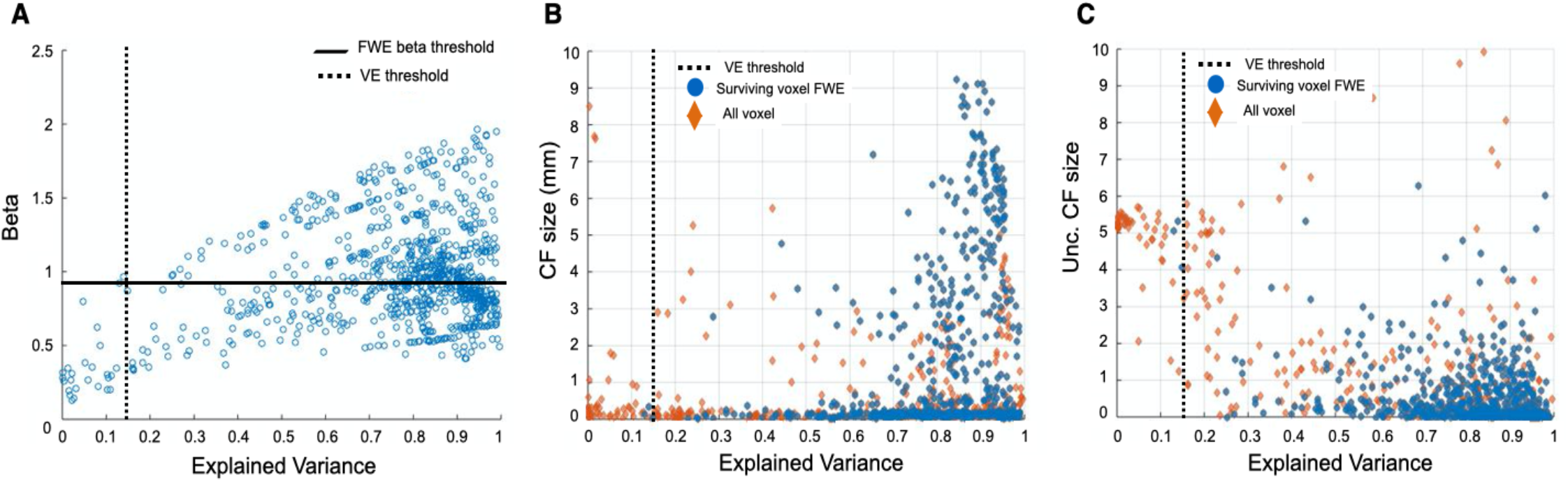
Comparison of thresholding approaches on a single subject level in V1>V2 area using RS1 data. In Panel **A,** the FWE *beta*-corrected thresholds obtained with both 95% CI and the standard VE of the model are shown. A direct comparison between the FWE threshold and the standard VE is presented in Panel B and C. Since we are interested in testing this FWE-corrected threshold in different conditions, we included voxels with CF size ~0 which are discarded. In Panel **B**, the relation between VE and the MCMC CF size is presented for all the voxels (orange diamonds). In blue the voxels surviving the 95% CI FWE beta-threshold are indicated. The standard VE threshold is not applied but indicated by a black dotted line. In Panel **C**, the relation between VE and the uncertainty associated with the CF size is presented.

This threshold was then used to compare the relation between RS-based CF size and VFM-derived eccentricity. Figure 5 shows that RS-based CF size does not increase with eccentricity within the early visual areas. While it is possible to notice an increase of CF size values with eccentricity only for the later visual areas (LO1 and LO2), especially in RS2. However, no significant differences were found between RS1 and RS2 scans in areas along the visual hierarchy.

**Figure 5.**
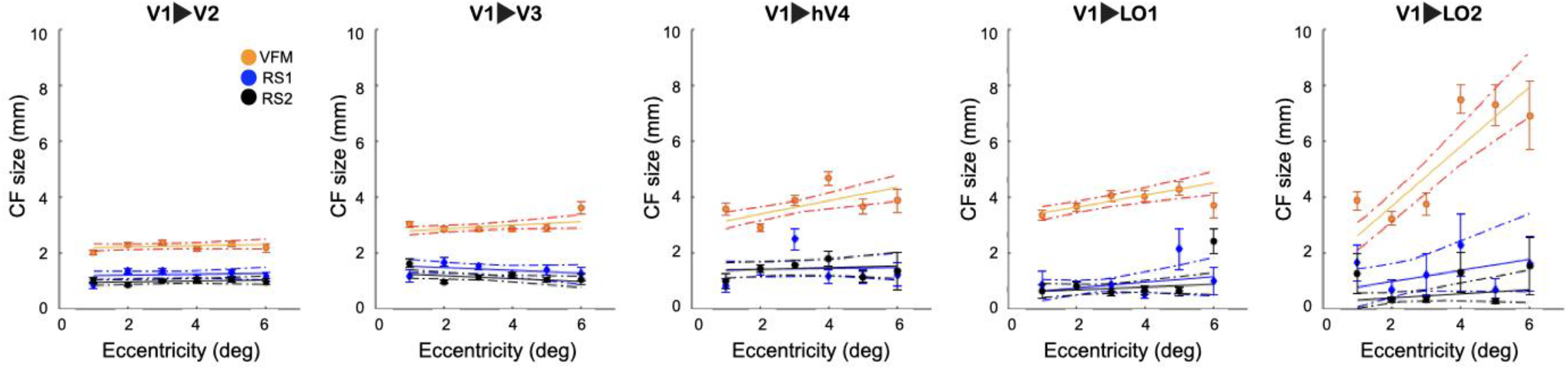
Connective field size as a function of VFM-based eccentricity for RS and VFM scans. For both RS and VFM scans, eccentricity was binned in intervals of 1 deg and a linear fit was applied. The average CF size was calculated for only voxels that survived the FWE 95%CI threshold. Each dot indicates the mean of the CF size for each bin. While the dashed lines correspond to the 95% bootstrap confidence interval of the linear fit. The VFM data is included in the figure as reference.

## 4. Discussion

In this study, we show that 3T RS-fMRI data is suitable for estimating local functional connectivity between visual cortical areas. Furthermore, we observed a good level of agreement between the standard and Bayesian (MCMC) CF models. This indicates that also the latter tool is suitable for studying the cortico-cortical properties of brains at rest. The obtained CF estimates are qualitatively similar to those previously observed for 7T RS-fMRI data. This further supports that sensitive estimations and associated uncertainties can be derived from 3T RS-fMRI data. Finally, we show that a FWE-corrected threshold based can be used as a complementary threshold to the standard V E to increase the reliability of estimates. Below, we discuss our findings in more detail.

### 4.1 The Bayesian CF framework applied to resting-state fMRI data

A good level of agreement was found between the CF and Bayesian CF maps estimated from RS and those estimated based on VFM (Figure 1 and table 1). As in the previous study from *Gravel et al. (2014)*, we observed substantial variability in the standard CF and Bayesian CF parameters obtained for different RS scans. Nevertheless, no significant difference was found between the two RS scans for any CF parameter. Even though the RS2 maps revealed a visuotopic organization that agrees better with the VFM-based maps (Figure 1, Panel A and D). In this respect, we note that RS2 was acquired after the visual stimulation (VFM).

Next, we investigated the characteristics of the uncertainty information. First, we checked whether a structure could be observed in the spatial distribution of this parameter. In order to do so, we compared the CF size, effect size (beta) and the associated uncertainties between the different conditions to determine if factors, like large-scale network interactions, physiological processes or measurement noise could possibly influence the uncertainties (Figure 3; Table 2). Similar to what observed on VFM data, no clear visuotopic organization was observed in the uncertainty-based maps obtained using RS data (Figure 3).

Second, we investigated (co)dependencies between the parameter estimate and the corresponding uncertainty. Similar to what observed in VFM data (Invernizzi *et al.*, 2020), a weak correlation was found between the Bayesian CF parameters (CF size and beta) and the corresponding uncertainties (correlation < 0.25) on RS data. Therefore as in VFM data, uncertainties can be treated as an additional, independent CF parameter for RS state data.

Finally, we compared the Estimated Variance (VE) of the model *versus* the FWE-corrected *beta* thresholding techniques based on the posterior distributions of the effect size (*β*). Based on the previous literature, we selected the cut-off value of the 95th percentile (Bornmann, 2013) as the FWE-corrected *beta* threshold. As previously shown by Invernizzi et al., *beta* thresholding is a reasonable approach for VFM data. Here, we show that the same approach, a FWE-corrected *beta* threshold based on the 95% CI is similarly valid for RS data (Figure 5 and 2S). Even though RS data is inherently more noisy than VFM data. We conclude that the FWE *beta*-threshold is a generally valid complementary approach to the standard VE for both VFM and RS data.

### 4.2 Connective Field estimations at 3T versus 7T

The CF method was previously used to reveal relevant aspects of resting-state brain activity using high-resolution 7T-fMRI. Crucially, in this study, we have extended the CF approach and assessed its performance at a lower-field strength (3T-fMRI). Higher magnetic fields can increase the signal-to-noise ratio, the tissue specificity and the spatial resolution of fMRI recordings. However, 3T scanners are much more abundant and more often used in clinical research than 7T ones. Here, we show that meaningful measures of cortical connectivity can be obtained based on 3T. We find that the CF maps obtained at 3T are in fair agreement to those obtained at 7T. RS-derived CF maps partly reflect the functional topographic organization revealed with pRF mapping –– regardless of the lower spatial resolution and signal-to-noise ratio of the 3T compared to 7T data. While higher magnetic field strengths allow for an enhanced spatiotemporal resolution, the temporal resolution of fMRI is limited by the hemodynamic response to neuronal activity, not by the magnetic field strength. This suggests that the spatially-weighted temporal correlations, captured by the CF method, suffices to reveal the underlying retinotopically organized connectivity between areas – albeit with somewhat lower spatial resolution and accuracy.

Examining the relationship between CF size and pRF eccentricity revealed a moderately positive correlation between VFM-derived parameters only, with increased CF sizes at higher pRF eccentricities (Fig. S3, supplementary material). This was most pronounced for the higher-order visual area LO2. In contrast, RS-derived CF size did not increase with eccentricity, neither within individual areas nor throughout the visual hierarchy (Figure 2 and Figure 5). The same trend was observed in the results obtained at 7T (Gravel *et al.*, 2014). These results might suggest a fluctuating signal-to-noise level in RS compared to VFM-data. Alternatively, it may indicate a lack of visual preprocessing in the peripheral visual areas when investigated with RS-data.

Our findings indicate that, despite the limited resolution of metabolism-sensitive measurements such as fMRI for determining the contribution of neuronal activity to hemodynamic signals, it is still possible to study the aggregate neuronal population properties at 3T using CF approaches.

### 4.3 Relation between resting state and functional architecture

Recent studies have shown that indirect measures of intrinsic neuronal activity, such as spontaneous BOLD fluctuations recorded during resting state, can still reflect the neuroanatomical connectivity organization that characterizes early visual cortical areas. These studies have allowed the assessment of both fine-grained within- and between-area interactions. This observed spatial specificity in spontaneous BOLD fluctuations can only emerge if these are anchored in the topographically organized architecture of the visual system as has been shown on multiple occasions (Biswal *et al.*, 1995; Baseler, Morland and Wandell, 1999; Azzopardi and Cowey, 2001; Haak *et al.*, 2013a). However, the neuronal and physiological basis of these BOLD patterns is still unclear. Whether spontaneous fMRI activity reflects the consequences of local population spiking activity, sub-threshold neuronal activity (Logothetis *et al.*, 2001), or metabolic relationships between neurons and astrocytes (e.g. neuro-vascular coupling) is still a matter of debate (O’Herron *et al.*, 2016; Pang *et al.*, 2017). It is possible that retinotopycally organized inter-areal BOLD coupling patterns reflect intrinsic activity in distant cortical areas, sharing similar selectivity in visual field positions. Likely due to “hard wired”, i.e. white bundles, coupling. Alternatively, these patterns may reflect the footprint of slow fluctuations that traverse the brain like “waves” (Logothetis and Wandell, 2004; Carandini *et al.*, 2015). Recent studies have unified these contrasting views by showing that both global fluctuations, in the form of propagating hemodynamic waves, and transient local coactivations are necessary for setting the spatial structure of hemodynamic functional connectivity (Pisauro *et al.*, 2013; Matsui, Murakami and Ohki, 2016). Taken together, these studies point to the multiple roles that neuroanatomical, physiological and vascular factors play in shaping spontaneous RS activity in a way that gives rise to visuotopically organized fluctuations in the BOLD signal. The similar visual field position selectivity revealed by RS- and VFM-derived CF maps, suggest a shared neuroanatomical origin.

### 4.4 Limitations and future directions

Here we compared the results of two data sets obtained from MRI measurements at different magnetic field strengths in different participants. Note that the results obtained using 3T and 7T dataset are consistent between participants for both standard and Bayesian CF models. This indicates the use of two different dataset is a valid approach to assess the CF models performance at different magnetic field strengths. However, for a more straightforward comparison, 3T and 7T derived results should ideally be obtained in the same participants.

The stimulus-agnostic character of the CF analysis invites applying the present approach also in other cortical regions, such as those involved in auditory, somatosensory, or motor processing.

Furthermore, future studies could investigate the correlations in the temporal and spatial domain in the cortex extending the Bayesian CF model to capture distinct dynamics in functional connectivity, and their relationship to different cognitive and behavioral states, both in health and disease.

## 5. Conclusion

We have shown that standard CF and Bayesian CF models are suitable tools to study the local functional connectivity in the visual cortical areas during resting state at 3T. Their output can be used to obtain quantitative estimates of the underlying neuronal architecture of the visual cortical areas. Both CF models provide results qualitatively similar to those previously observed at 7T. Using our novel Bayesian CF modeling approach, we show that new independent CF parameters can be derived: specifically, uncertainty and effect size. These parameters provide a novel way of assessing the statistical significance of our modeling approaches, namely a threshold based on effect size, which is an alternative approach to the classical variance explained and are necessary when comparing conditions, models and/or groups.

### Supplementary Material

#### Bayesian connective field model: MCMC CF option B

The following description is adapted from Invernizzi et al. (2020).

Based on the CF definition used in the standard approach (Haak *et al.*, 2013b), a linear spatiotemporal model and a 2D symmetric Gaussian connective field model (2) are used to create a predicted time serie (*p*(*t*)) which is fitted to the time series *y*(*t*)of a target location (1).

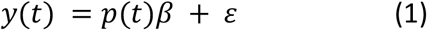

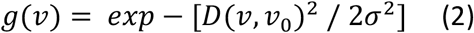

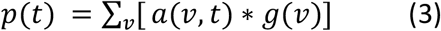

Where the predicted fMRI signal *p*(*t*) is obtained by the overlap between the CF model*g*(*v*) and the neuronal population inputs *a*(*v*, *t*), that are defined as the BOLD time series (converted to percent signal change) for voxels (*v*) (see eq. 3). In equation 1, *β*defines the effect size and *ε*is the error term. The 2D symmetric Gaussian CF model of voxel (*v*),*g*(*v*)is defined based on the shortest three-dimensional distance *D*(*v*, *v*_0_) between a voxel (*v*) and the proposed CF center (*v*_0_) on a triangular mesh representation and σ, which defines the width of the CF. *D* is computed using Dijkstra’s algorithm (Dijkstra, 1959) whileσis constrained to the range[*r*_0_ *r*]using a latent variable*l*_σ_(Zeidman *et al.*, 2018). A flat prior is assumed for σ. Therefore, the prior for the latent variable*l*_σ_ is defined as a normal distribution *N* 0,1 (see equation 4). As explained in Zeidman et al.(Zeidman *et al.*, 2018), each latent variable is assigned to a prior distribution that represents our beliefs for that CF parameter, before the model fitting.

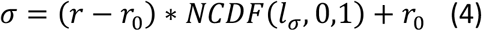

Where*r*is the maximum radius and *r*_0_is the smallest allowed radius for the CF width - that can be an arbitrarily small non-zero number, which here were set to 10.5° and 0.01°, respectively. *NCDF*indicates the normal cumulative distribution function.

The MCMC is an iterative sampling approach. During each iteration the parameters for a new CF are set and the fit is compared against the current one. A new location will be selected using the distance to the current position (*d*_*current*_). Based on the distance matrix (D), the maximum step (*ms*) possible in the source region was defined as half the maximal distance from the current position (*d*_*current*_) (5). Latent variable *l_S_*, is randomly drawn from a normal distribution N(0,1) which results in a flat prior for the step size (*step*) between 0 and the maximum step [0 *ms*] (5, 6). The updated sampling position (*v*_0 *proposal*_) is defined as that position for which the distance to the current position is as close as possible to *step*. If multiple locations are found, only one is drawn randomly.

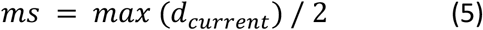

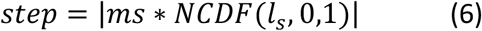

Note that for the first iteration the CF center (*v*_0_) was randomly selected from the source region.

Simultaneous with an updated sample location, an updated width for the CF is calculated. The *l*_σ *proposal*_ is drawn from a gaussian distribution centered around the current value with a width *w*_*proposal*_ (7).

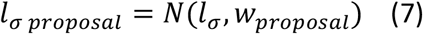

The effect size (*β*) is estimated in parallel to the other CF parameters and constrained to be positive (Zeidman *et al.*, 2018) using the following equation:

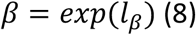

A latent variable *l*_*β*_ was defined with a prior distribution *N* −2,5 and the next*β*value was controlled by *l_β proposal_* (9).

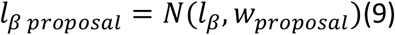

In this study, the initial values of*l*_σ_, *l*_*β*_and *w*_*proposal*_were set to 1, −5 and 2, respectively.

At each iteration of the MCMC, the updated CF parameters (σ, *β*) were estimated using the following steps. First a predicted fMRI signal*p*(*t*) is generated from the source region using eq 2. Note that *g*(*t*) was scaled to ensure that the total area under the gaussian, as calculated across the full source region, was equal to one. Second, the error per time point *e*_*t*_between the measured fMRI signal (*y*(*t*)) and the predicted fMRI signal *p t* was calculated.*e*_*t*_is calculated via subtraction of the predicted signal *p*(*t*) from the measured fMRI signal. Then, the log-likelihood*L*_*t*_ associated with*e*_*t*_was estimated using equation (10). We assumed that *e*_*t*_follows a standard normal distribution: *N*(0,1). After estimating the mean and standard deviation of ∈ (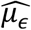 and 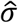), we calculated the maximum likelihood estimates (*MLE*, eq. 11).

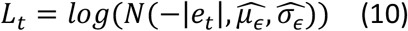

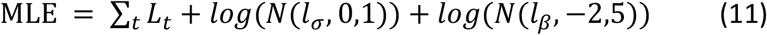

At this point, MLE of the proposal iteration is compared to the last accepted (current) sample based on an Accepted ratio score *Ar* (12).

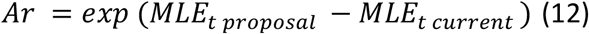

*Ar*was compared to a pseudo-random acceptance score defined as a normal distribution *N* 0,1 and only if the *Ar* was higher, the respective latent variables were updated. Based on the accepted *l*_σ_, *l_S_* and *l*_*β*_ values, a new CF was defined and a new MCMC iteration took place.

## Acknowledgements

We want to thank Barbara Nordhjem for data collection and Hinke N. Halbertsma for data collection and preprocessing (manual segmentation and ROI definitions).

## Supplementary Figures

To check the possible influence of the ICA-AROMA denoise procedure, the same quantification analysis was computed on non-denoised RS data. Similar maps (Figure 1S) and correlation values (Table 1S) were observed indicating that the ICA-AROMA denoise procedure on RS-fMRI data did not influence the final CF outcomes.

**Figure 1S.**
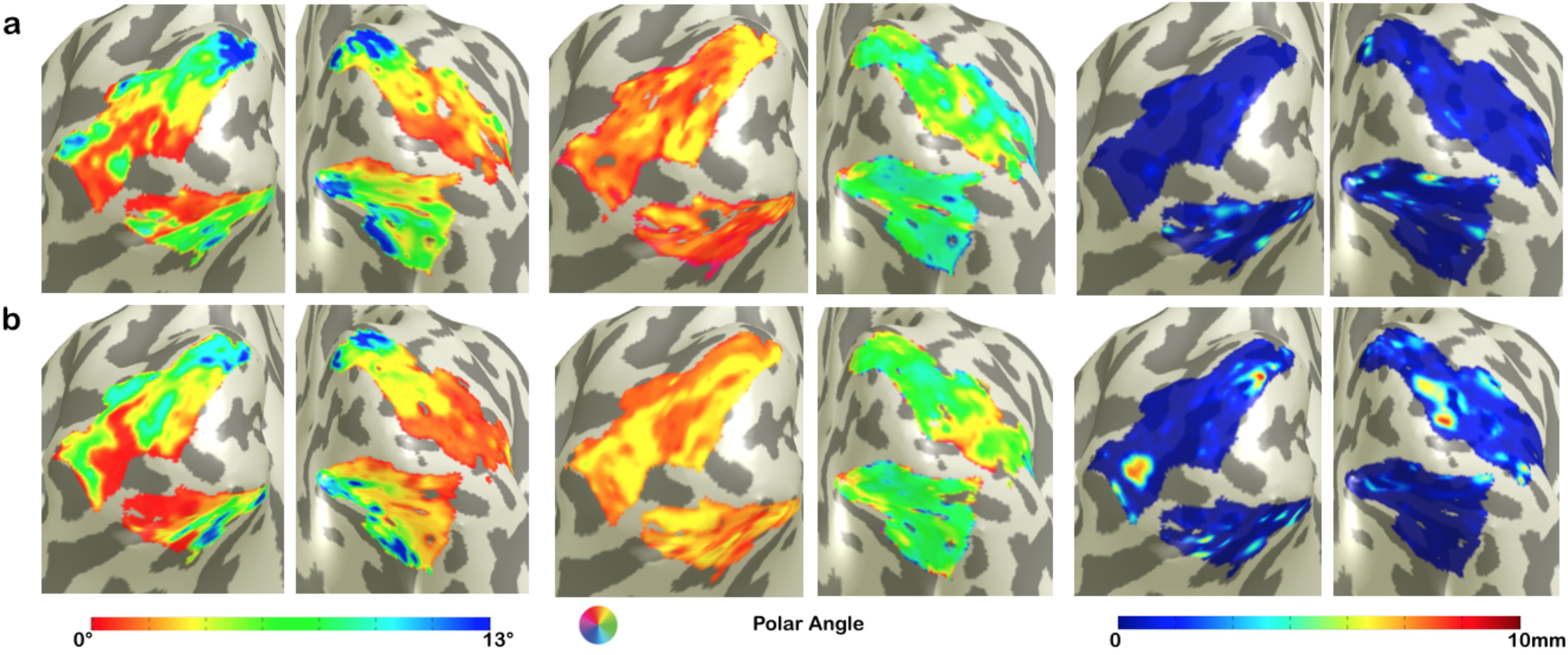
Visualization of CF maps of not-denoised RS data for a single subject. From left to right: eccentricity, polar angle and CF size. Panels A and B show CF parameters for each RS run before applying ICA-AROMA denoising procedure.

**Table. 1S.**
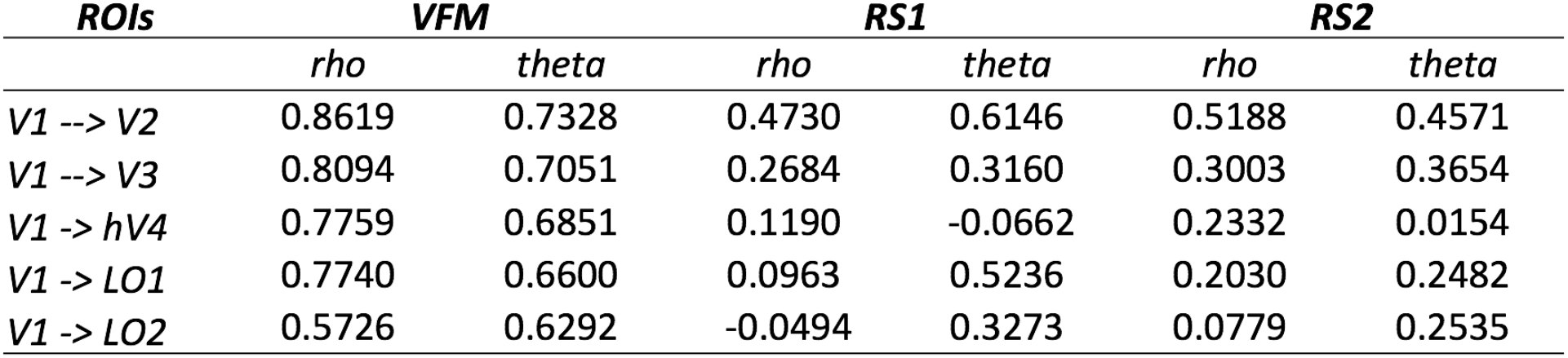
Correlation between Bayes pRF and CF maps obtained from VFM and no-denoised RS data at group level. To estimate and compare the level of agreement of CF maps, we computed the correlations between eccentricity (*rho*) and polar angle (*theta*) parameters obtained using standard CF models to the Bayesian pRF *rho* and *theta (*gold standard).

**Figure 2S.**
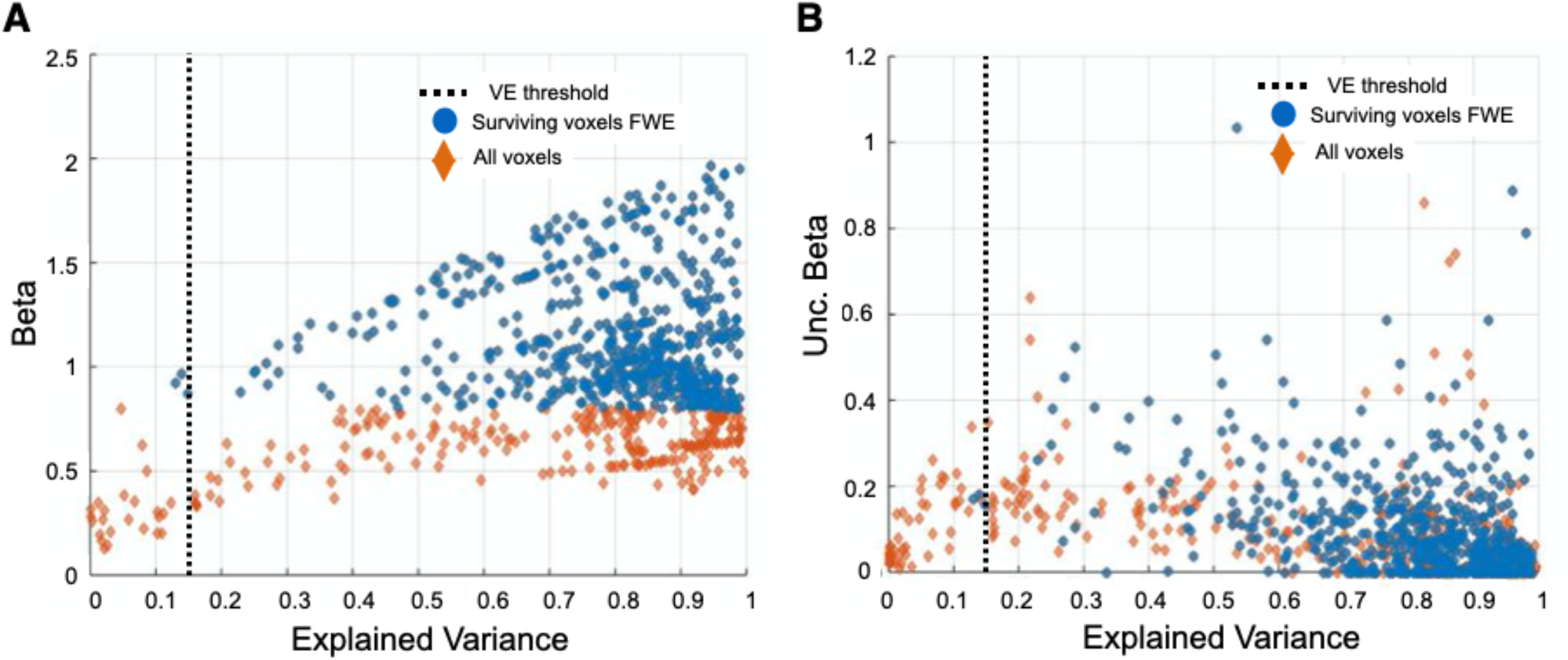
Comparison of thresholding approaches on a single subject level in V1>V2 area using RS1 data. In Panel **A**, the relation between VE and the beta parameter is presented for all the voxels (orange diamonds) and only for the ones surviving the 95% CI FWE beta-threshold (blue dots). The standard VE threshold is not applied but indicated by a black dotted line. While in Panel **B**, the relation between VE and the uncertainty associated with beta is presented.

**Figure 3S.**
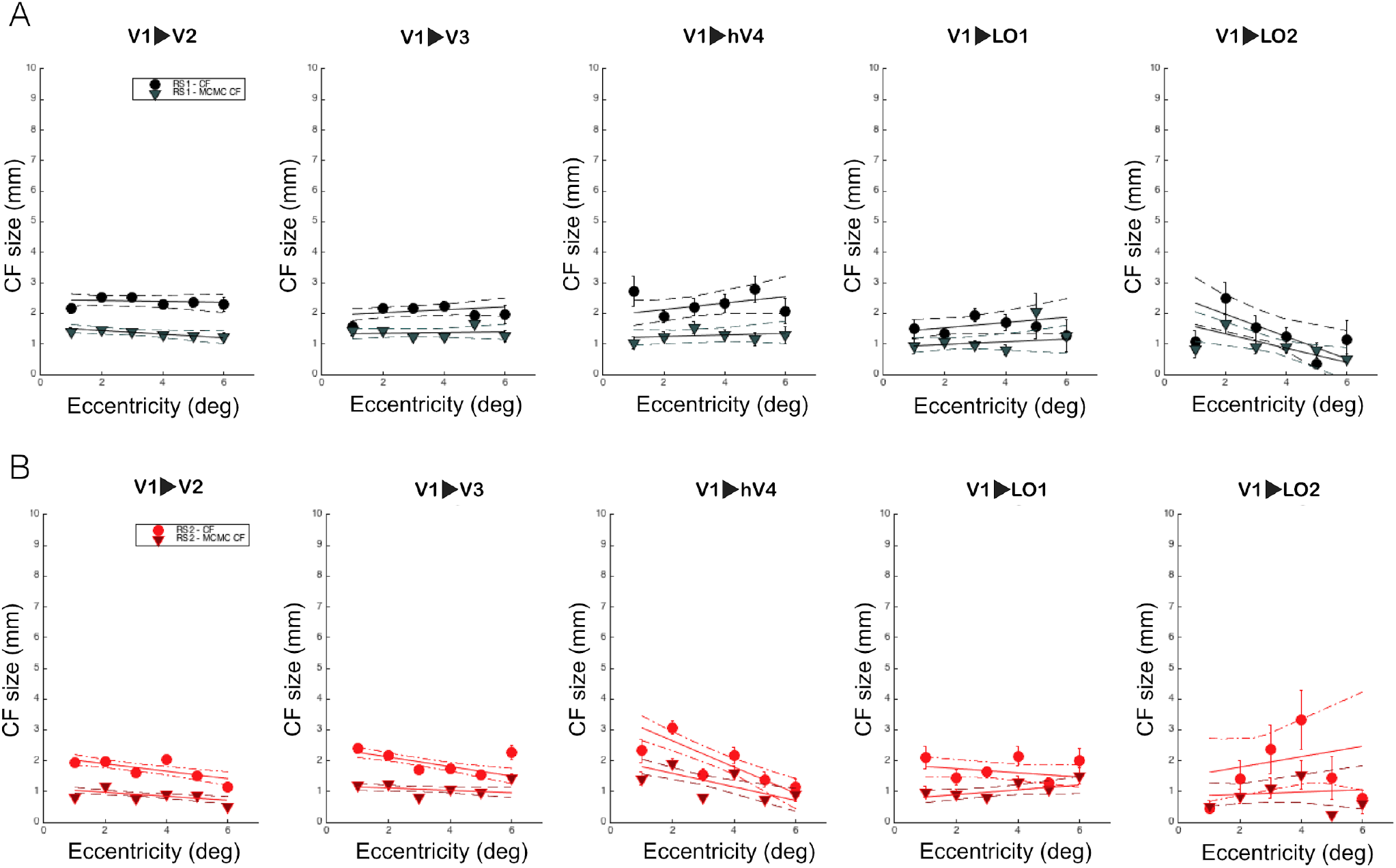
Connective field size as a function of pRF eccentricity for RS scans. For standard and MCMC CF models, eccentricity was binned in intervals of 1 deg and a linear fit was applied. The CF size was initially weighted with variance explained higher than 0.15. Each dot and triangle indicate the mean of the CF size for each bin. While the dashed lines correspond to the 95% bootstrap confidence interval of the linear fit. In Panel **A**, CF models were applied to RS1 scan while, in panel **B** to RS2 scan.

